# A model for pH coupling of the SARS-CoV-2 spike protein open/closed equilibrium

**DOI:** 10.1101/2020.10.31.363176

**Authors:** Jim Warwicker

## Abstract

SARS-CoV-2, causative agent of the COVID-19 pandemic, is thought to release its RNA genome at either the cell surface or within endosomes, the balance being dependent on spike protein stability, and the complement of receptors, co-receptors and proteases. To investigate possible mediators of pH-dependence, pKa calculations have been made on a set of structures for spike protein ectodomain and fragments from SARS-CoV-2 and other coronaviruses. Dominating a heat map of the aggregated predictions, 3 histidine residues in S2 are consistently predicted as destabilising in pre-fusion (all 3) and post-fusion (2 of 3) structures. Other predicted features include the more moderate energetics of surface salt-bridge interactions, and sidechain-mainchain interactions. Two aspartic acid residues in partially buried salt-bridges (D290 – R273 and R355 – D398) have pKas that are calculated to be elevated and destabilising in more open forms of the spike trimer. These aspartic acids are most stabilised in a tightly closed conformation that has been observed when linoleic acid is bound, and which also affects the interactions of D614. The D614G mutation is known to modulate the balance of closed to open trimer. It is suggested that D398 in particular contributes to a pH-dependence of the open/closed equilibrium, potentially coupled to the effects of linoleic acid binding and D614G mutation, and possibly also A570D mutation. These observations are discussed in the context of SARS-CoV-2 infection, mutagenesis studies, and other human coronaviruses.

## Introduction

Viruses must deliver their genome to the host cell, and for membrane enveloped viruses this occurs either at the cell surface, and fusion with the plasma membrane, or through fusion with an intracellular membrane, subsequent to import into the cell. Coronaviruses are membrane enveloped viruses that deliver their RNA genome either through fusion at the plasma or endosomal membranes [1]. Factors that determine which fusion route dominates for a particular virus include cell-surface receptor and availability (dependent on cell type), co-receptors, spike (S) protein structural stability, and which proteases carry out the S1/S2 and S2’ cleavages [2]. For SARS-CoV-2, causative agent of the world-wide COVID-19 pandemic, cell surface ACE2 is the primary receptor [3], a novel protease specificity has evolved [4], and the primary route of infection (although this depends on protease location and other factors) may be endosomal [5]. Considering pH values in early (6.3) and late (< 6) endosomes, with regard to virus entry and genome release, and also the export secretory pathway for newly assembled viruses (pH 5.5 in secretory vesicles), it is clear that environmental pH is likely to play a part in the infection process [6]. The primary focus for molecular response to pH is the coronavirus spike protein, with analogy to the role of influenza hemagglutinin, where pH-dependent conformational changes underpin endosome-mediated infection [7].

Spike glycoprotein structures for SARS-CoV-2 are accumulating rapidly, mostly in two broad categories, cryo-electron microscopy (cryo-EM) analyses of the S glycoprotein soluble ectodomain (S protein for brevity), and X-ray crystallographic structures of fragments, largely the receptor binding domain (RBD) [8, 9]. In both categories, structures may be S protein alone (normally as a trimer in studies of full length protein) or in complex with receptor (ACE2) or antibody fragments. Two large-scale structural themes are apparent. First, pre-fusion, the RBD populates open (up) or closed (down) conformations, with the open form competent for ACE2 binding. Second, comparison of pre-fusion (S1 and S2) and post-fusion (S2 only) structures reveals not only the absence of the shed S1 protein, but also the large conformational change that uncovers the putative fusion peptide, moving it towards the anticipated location of the target cellular membrane [10]. Biophysical and thermodynamic data for SARS-CoV-2 S protein are accumulating more slowly than the structural database. Deep mutational scanning of the effect that amino acid substitutions make on RBD expression and ACE2 binding has been performed in a yeast display system [11]. This study is targeted at informing which regions are likely to be more evolutionary restricted, for vaccine design, but it is also valuable for assessing the structural and functional importance of residues for which computational work provides predictions.

Protein regions that mediate biological pH-dependence can be predicted with pKa calculations [12]. The pH-dependence of free energy difference between folded and unfolded states is directly related to the difference in protonation state (charge) between folded and unfolded states, and thus the pKas [13]. Continuum electrostatics has been widely used for pKa prediction [14], and constant-pH molecular dynamics allows the explicit inclusion of conformational variation into predictions [15], particularly where a structural focus has already been established. SARS-CoV-2 full length S protein is large (1273 amino acids), locations of sites that may determine pH-dependence have not been determined in detail, and there are an increasing number of cryo-EM structures covering various extents of open and closed conformations in the trimer, at resolutions typically around 3 to 4 Å. In this scenario, a reasonable strategy is to apply continuum electrostatics calculations over the available structures and seek consensus results as the basis for prediction. The algorithm used for this study has been reported previously [16], as has a web implementation of a reduced version [17] that benchmarked the method against known pH-determinants for influenza hemagglutinin. That study also identified two histidine residues predicted to be buried and destabilised in both pre-fusion and post-fusion SARS-CoV-2 S protein. Reported currently is an extensive study of calculated ionisable group pKas over the rapidly expanding dataset of S protein structures, for which averaging reveals features that are predicted to stabilise or destabilise the S trimer. The results are further studied to reveal groups that could couple changes in pH to structural stability, and therefore with a predicted impact on the viral life cycle.

## Results and discussion

### Predicted electrostatic frustration in the S protein

The set of 24 SARS-CoV-2 S trimer cryo-EM structures (72 monomers) was assembled from the PDB/RCSB [8] in August 2020. An offline version of the code in use at www.protein-sol.manchester.ac.uk/pka [17] was used to predict pKas, with each S trimer covered in a single calculation. Predicted pKas were assembled into a heat map, split according to S1 and S2 pre-fusion proteins and S2 in the post-fusion form, and sorted according to up and down (RBD) monomer instances. Additionally, pKas were averaged for open and closed monomers and displayed in the same colour-coded format as the web application. Thus, red denotes destabilisation (electrostatic frustration in the fold) and blue stabilisation, in both heat map (Figure 1, Supplementary Table) and molecular view (Figure 2).

**Figure 1.**
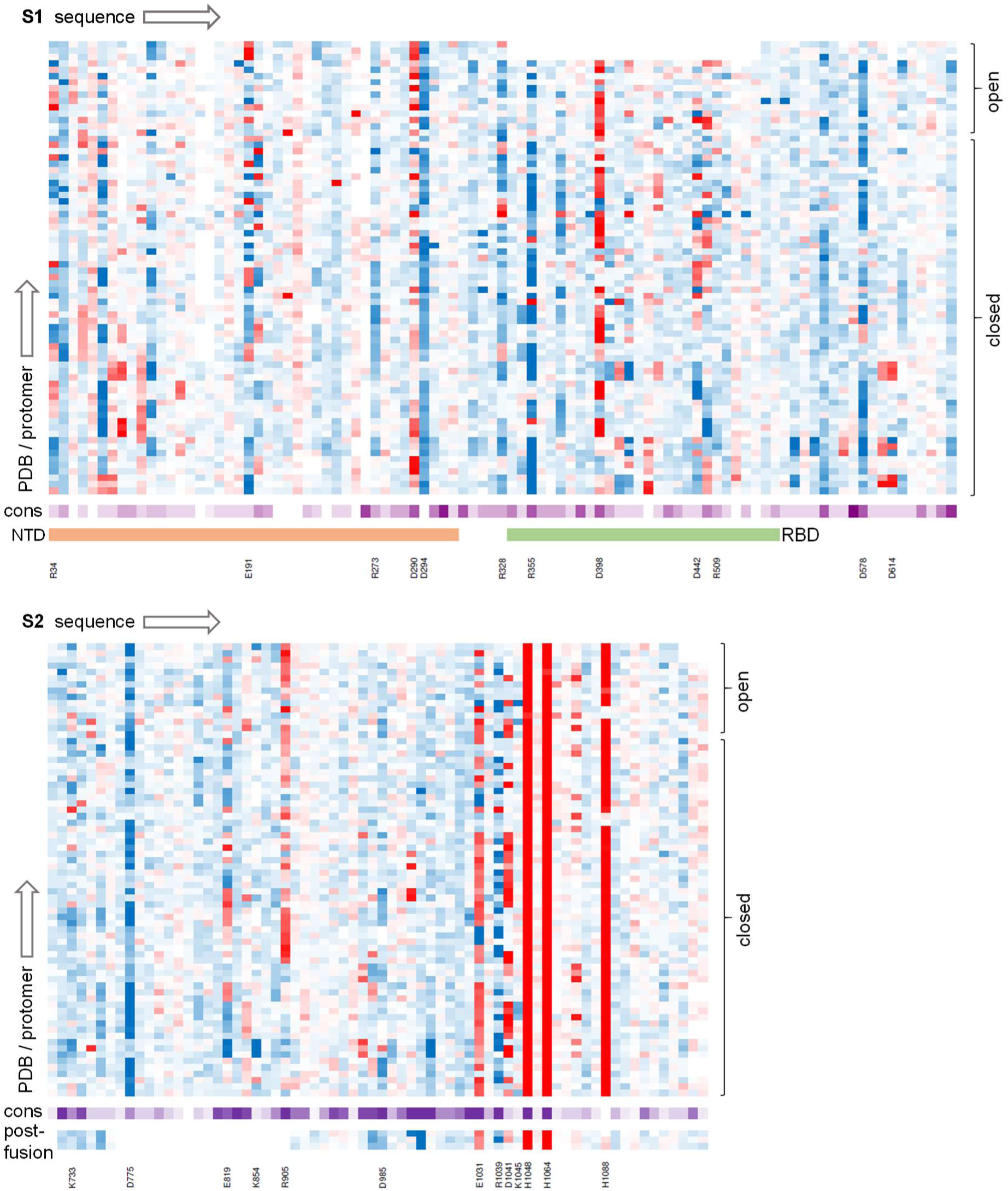
Ionisable group properties in the trimer dataset. Predicted changes in ionisable group pKas in a heat map, from brighter red (destabilising) to darker blue (stabilising), with separate panels for S1 (upper) and S2 (lower) proteins. Calculations are grouped according to open and closed monomers (RBD), as indicated. Sequence conservation across coronaviruses is shown with a deeper purple for greater conservation. Residues discussed in the text are listed, and the extent of NTD and RBD domains is displayed for S1.

**Figure 2.**
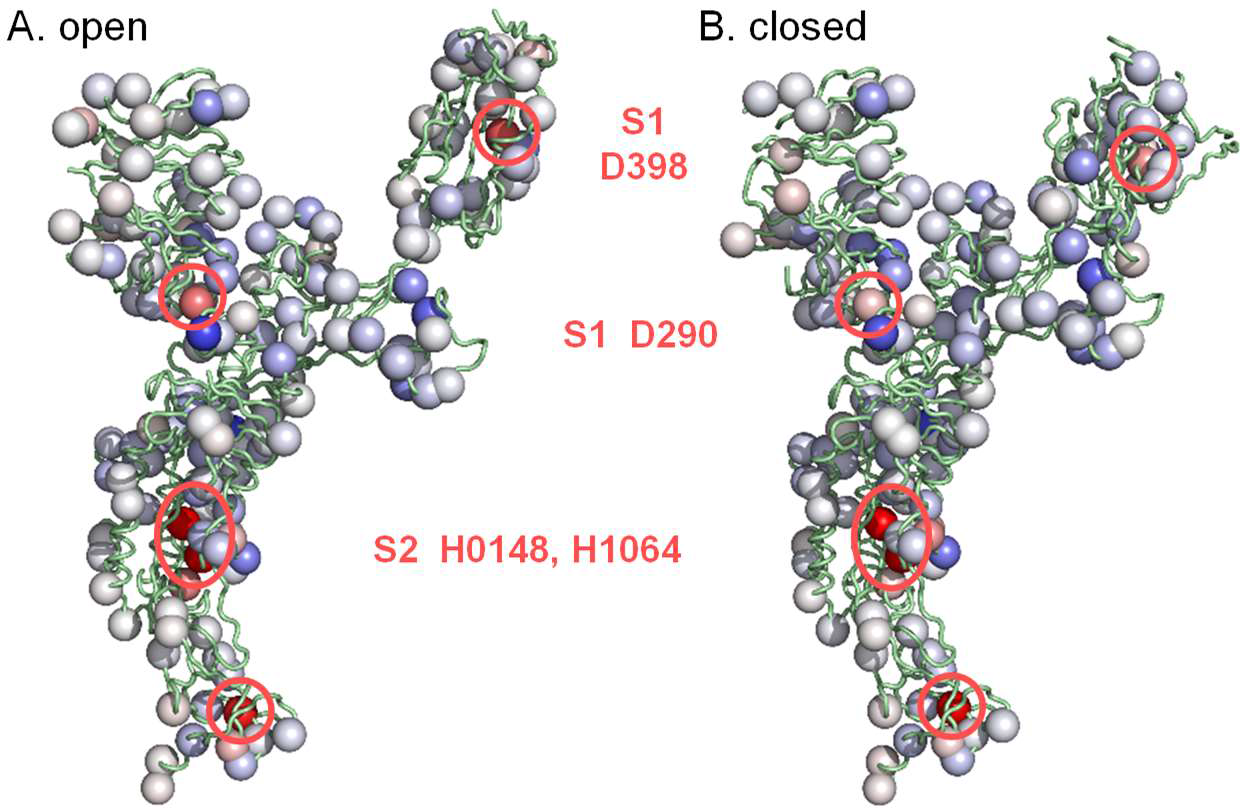
Averaged predicted ΔpKas in molecular display. Ionisable groups are coloured using the same scale as Figure 1 (red is destabilising, blue is stabilising), here with averages over the open (A) and closed (B) form monomers. Sites of greater destabilisation are highlighted and their identity noted.

Most obvious in Figures 1 and 2 are three histidines in S2 of the pre-fusion protein, two of which (H1048 and H1064) are conserved in coronaviruses and were reported previously [17]. Histidines 1048 and 1064 are buried in both pre- and post-fusion S2, in a subdomain that adopts the same fold in pre- and post-fusion structures [10]. It was suggested that the electrostatic frustration exhibited by these residues would be relieved if the subdomain transiently unfolded during the transition to post-fusion S2 [17]. A third histidine in S2 (H1088) is also destabilised consistently in the pre-fusion structure, but is present only in SARS-CoV-2, of the 10 aligned sequences (Figure 1), and is not buried or destabilised to the same extent in the post-fusion structure (6xra) [10]. Histidine 1088 is predicted to contribute to stabilisation of the post-fusion protein relative to the pre-fusion protein at acidic pH values, such as endosomal pH, where histidine would normally be ionised.

There are no other ionisable groups in the trimer dataset predicted to possess the extent and consistent destabilisation of the 3 buried histidines already noted. Rather, there is a mix of groups in differing environments, including partially buried charge pairs, charge-dipole interactions, and charge pairs at monomer-monomer interfaces and surfaces, examples of which are noted in Figure 1, and discussed here. Amongst these are the charge pairs E1031 - R1039 and D1041 - K1045, of which the former is relatively conserved amongst coronaviruses and the latter less so. These charge interactions are partially buried, and thus relatively strong, and in both pre-fusion and post-fusion structures are adjacent to monomer-monomer interfaces, in the same domain as the buried H1048 and H1064.

Considering amino acids not close to the buried histidines, D985 is an interesting case, positioned to act as an N-terminal helix cap in a region that has been engineered to incorporate a stabilising double proline substitution [18]. The 985 position is conserved as either aspartic or glutamic acid in coronaviruses, consistent with it being an important site for spike protein stability. Aspartic acid 294 is another N-terminal helix cap, leading to consistent predicted stabilisation, although it is not conserved across coronaviruses. Several charge pairs exist, with partial burial leading to increased interactions relative to purely solvent exposed location, but generally predicted to be stabilising, including R34 - E191, R328 - D578, D442 - R509, and K773 - D775. Residues E819 and R905 are each partially buried and form interactions with mainchain and sidechain hydrogen bonding groups, which are calculated to be stabilising in some monomers and destabilising in others.

The primary focus of this study is those groups that are predicted to be frustrated in S protein trimers, leading to pKa changes and the potential for pH-dependent effects. From Figure 2, this filter greatly narrows the ionisable groups of most interest. They are the buried histidines in S2, and two charge pairs in S1.

### Partially buried pair D398 - R355 is predicted to couple with the open/closed equilibrium

The two charge pairs that feature predicted electrostatic frustration, when averaged over the dataset, are D398 – R355 in the RBD, and D290 – R273 in the NTD (Figure 2). Focussing on D398 – R355, the aspartic acid has very little solvent accessibility in all structures studied, whilst the arginine burial varies according to structure (Figure 3). Average calculated D398 pKa for closed form monomers is 5.8, and for open form monomers 7.1, with 10 of 11 open monomer pKas at 7.0 or above, and the remaining one slightly lower than the intrinsic pKa of 4.0. In contrast, predicted D398 pKas for the closed form monomers vary across a range of stabilising and destabilising values. The environment of R355 changes between open and closed forms, forming a larger interface with the NTD of a neighbouring monomer in the closed form (Figure 3), but also with variation in the closed interfaces. Indeed, for the 58 instances of RBDs in the closed conformation, R355 solvent accessibility correlates with D398 pKa (r=0.55, p<0.00001). In closed form monomers, modulation of the strong electrostatic coupling between ionised D398 and R355 sidechains (which balances the D398 burial penalty) leads to the variation in predicted D398 pKa values. Greater occlusion of R355 at the interface gives increased coupling of ionised D398 and R355. The consequence of these calculations is a predicted contribution that destabilises the open form relative to closed from, with a pH-dependence that depends on D398 pKa.

**Figure 3.**
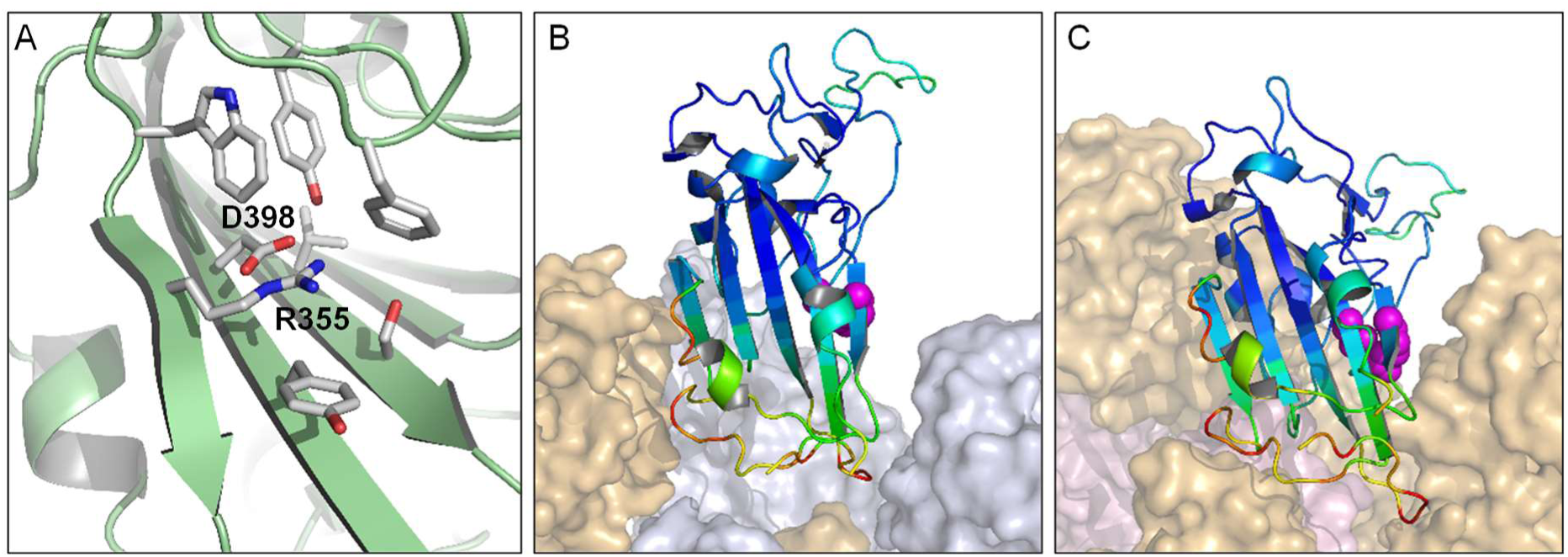
The environment of charge pair D398 – R355 in open and closed forms. A view of D398 – R355 (6lzgB) indicates the extensive burial of D398 and partial burial of R355 (A). The charge pair is represented as magenta spheres in the 6lzgB crystal structure, which is aligned to open (B) and closed (C) monomers in the 6vsb trimer context. Colouring of the RBD crystal structure shows a substantially higher crystal B-factor for a region on one side of the charge pair.

In order to test the calculations for cryo-EM structures, a set of X-ray crystal coordinate sets was assembled for the RBD of SARS-CoV-2 (crystallised alone or in complex with ACE2 or antibody fragments). Since these RBD structures are not within the context of the S trimer, they represent open conformations. The average predicted D398 pKa for the 10 RBD structures is 7.1, all substantially destabilising, with the lowest at 5.8 (6m0j chain E), in agreement with the cryo-EM structure calculations.

Crystal structures are available for RBDs of 4 of the 7 human coronaviruses (SARS-CoV-1, SARS-CoV-2, MERS, HCoV-HKU1), and a cryo-EM structure for the closed conformer of HCoV-OC43. For the remaining two human coronaviruses, HCoV-NL63 and HCoV-229E, D398 and R355 (SARS-CoV-2 numbering) are not present in the alignment (Figure 4). A characteristic of the charge couple in open monomer SARS-CoV-2 calculations is that ionised R355 is stabilised and ionised D398 destabilised. This is also evident for predictions in the RBDs of SARS-CoV-1 and MERS, but not for HKU1 or HuOC43 (Figure 4). The collapse of charge coupling between residues equivalent to D398 and R355, and the consequent loss of predicted pH-dependence, for HKU1 and HuOC43 are due to sequence and structure changes in this region. In HKU1 the residue equivalent to Y396 is valine, and in HuOC43 it is threonine, the smaller sidechains increasing solvent accessibility at the site corresponding to R355. The residue equivalent to F464 is glycine in HKU1, and this region of HuOC43 folds away from the charge couple, the effect in both HKU1 and HuOC43 being to reduce the solvent accessibility of residues equivalent to both R355 and D398. Thus, pH-dependent differential stabilisation of open and closed trimers is predicted for the severe human infections, SARS-CoV-1, SARS-CoV-2, and MERS, but not for the other 4 known human coronaviruses, HKU1, HuOC43, NL63, 229E.

**Figure 4.**
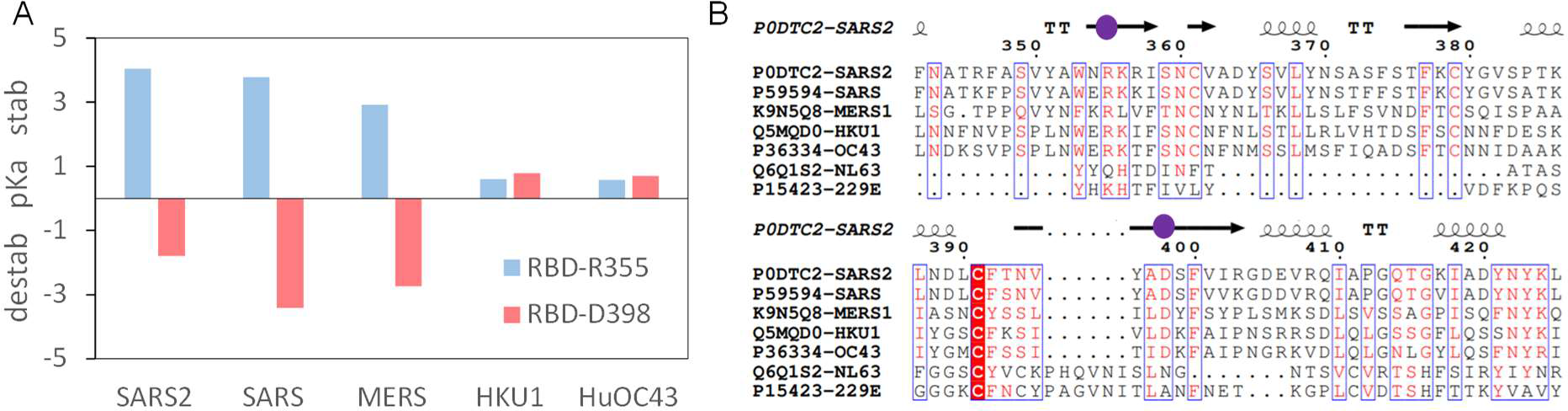
The D398 – R355 pair in human coronaviruses. Predicted destabilising and stabilising ΔpKas of R355 and D398 for the 5 human coronaviruses with available RBD structural information (A). Sequences for the 7 human coronavirus are aligned over this region (B), with secondary structure added from 6vsbB, and sequence conservation displayed (ESPript). Locations of R355 and D398 are indicated with purple spheres.

Charge pair D290 – R273 in the NTD is qualitatively similar to R355 - D398, with D290 having very low solvent accessibility and R273 greater solvent accessibility. Interpreting open versus closed is not as straightforward for the NTD, since the NTD most affected by an open monomer (RBD up) will be one of the other two monomers in the trimer. Thus, NTDs were re-assigned as open or closed according to the relevant contact RBD, giving an average predicted D290 pKa of 3.9 for 12 NTDs neighbouring an open RBD, and 5.5 for 57 other NTDs in the trimer dataset. The correlation observed for D398 – R355, between arginine solvent accessibility and aspartic acid pKa, is not seen here for D290 – R273. Given this lack of correlation and the lower predicted D290 pKas, compared with D398, the D290 – R273 pair is not pursued further in this report.

### Environment of the D398 - R355 charge pair

A number of reports impact on the prediction that D398 – R355 contributes to a pH-dependence of the open/closed equilibrium, with more open forms being increasingly disfavoured as pH rises through the mild acidic range. A report of S trimer structures at acidic pH, including calorimetric data, leads to the suggestion of a pH-dependent switch that plays a role in avoiding immune surveillance of RBD open forms of the S protein, and also that D614 is a part of the switch [19]. This is an intriguing hypothesis in the context of the establishment of the D614G mutation in SARS-CoV-2 genomes over the first year of the pandemic, and reports that the mutation may bias towards the (ACE2 binding) open form [20], altering barriers to conformational change between open and closed [21]. Interestingly, D614 is not prominent in pKa calculations with the trimer dataset collated here, with little indication that D614 has sufficiently strong interactions to generate a substantial pH-dependence. In very recent reports, an S trimer that binds linoleic acid (LA) has been studied [22]. The monomer – monomer interface, in regions adjacent to both the D398 – R355 charge pair and to D614, is more closely packed in the fatty acid bound structure than in the majority of closed S trimers (Figure 5). The LA binding region is in a subdomain of the RBD that lies adjacent to the D398 – R355 pair, and which consistently in crystal structures has higher B-factors than the subdomain responsible for ACE2 binding (Figure 3). If the flexibility of this region is dependent on fatty acid binding, then it could couple to alteration of R355 solvent accessibility and thus, according to the current work, D398 pKa and pH-dependent conformational biasing of closed versus open conformation. Further, the effects of LA binding extend also to the region around D614, where close packing leads to the hypothesis that D614 interactions act as a latch for the closed state which is lost with D614G mutation [20].

**Figure 5.**
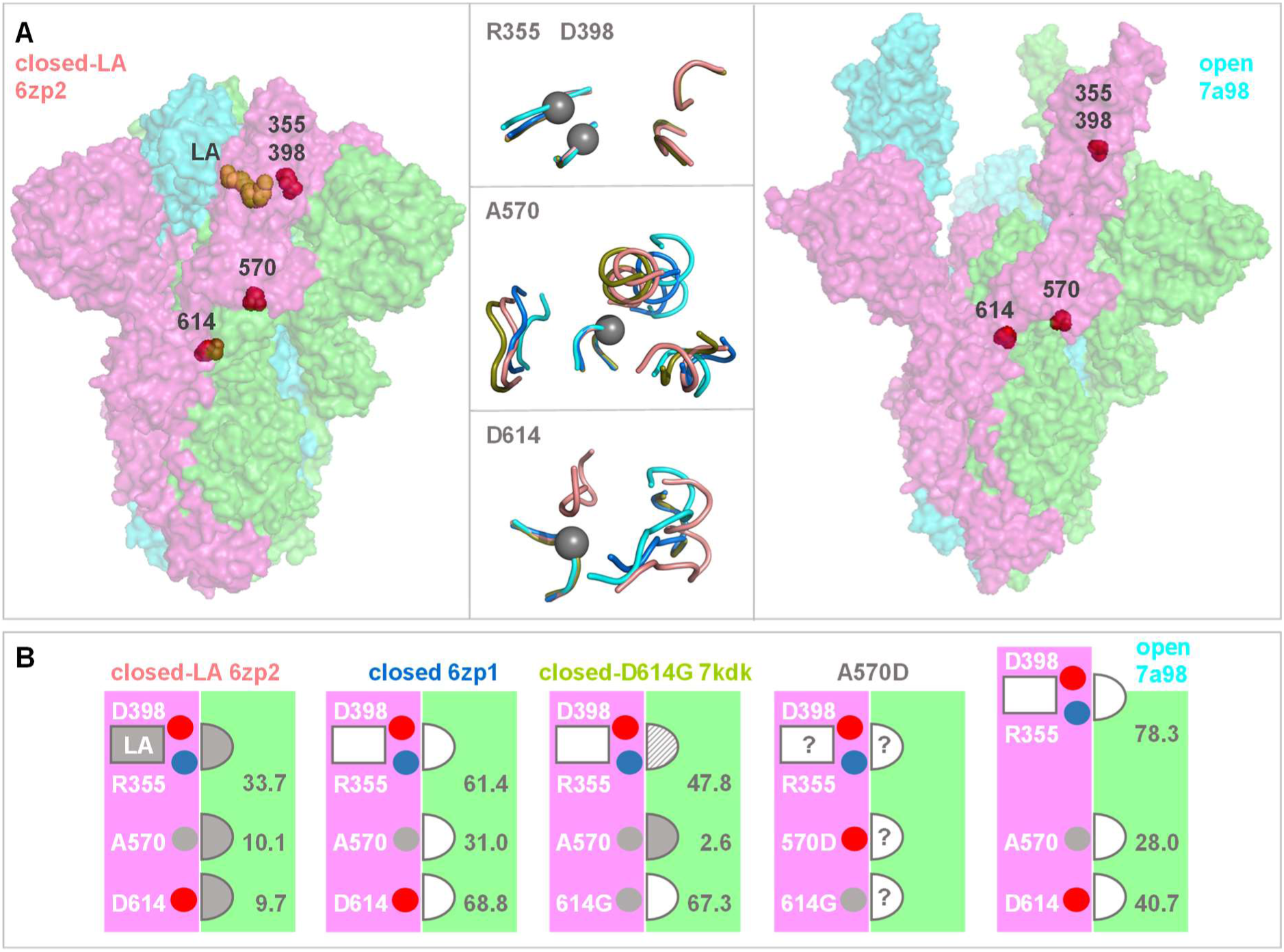
Structural comparisons of monomer – monomer interfacial regions in S protein trimers. (A) Spike protein ectodomain trimers are drawn in surface representation for a closed LA-bound structure (left, 6zp2), and a fully open trimer extracted from a complex with 3 receptors (right, chains A, B, C from 7a98). Sites of current interest within these transparent surfaces are indicated by red spheres, and LA is drawn in orange. In the horizontal centre are tube cartoon representations of 6p2 and 7a98, alongside 6zp1 (closed, no LA) and 7kdk (closed, D614G), drawn for regions within 10 Å of R355 (upper), A570 (centre) and D614 (lower). Centres are shown with grey spheres on the Cα atoms of D398/R355, A570, or D614, and other than the regions containing these residues, all other segments are from a neighbouring monomer. The cartoons are colour-coded according to structure, 6zp2/orange, 6zp1/marine, 7kdk/olive, 7a98/cyan. Separate structural alignments were made for each panel, aligning the peptides containing R355, A570, or D/G614 to that of 6zp2. (B) Structure colour-coding is reproduced in the titles of these schematic plots. Each schematic represents the accessibility of each of the R355, A570, D614 residues at the interface between neighbouring monomers (pink and green). LA is bound only to 6zp2, with the most close-packed interface as determined from panel A, and also from the listed solvent accessibility values (Å^2^). The semi-circular sites are indicated throughout as closely packed (filled in), not closely packed (unfilled), or intermediate (dashed). In the absence of a known structure for interfacial regions in an S trimer bearing the A570D mutation, the degree of packing at monomer - monomer sites is unknown, as shown. In the schematic images, residues of interest are indicated as small spheres, red/acidic, blue/basic, grey/aliphatic.

Environments around D398 – R355 and D614 are summarised in Figure 5A, for 6zp2 (closed S trimer with LA bound [23]), 6zp1 (closed S trimer without LA [23]), 7kdk (closed D614G S trimer [24]) and 7a98 (open S trimer [25]). In the LA-bound structure, the environment is more restricted for R355 and D614 (and solvent accessibility lower, Figure 5B), than for the closed trimer without LA, or for the open trimer. The D614G structure is more open around G614, and intermediate (between closed trimers with and without LA) around R355. Comparison of LA bound and unbound closed trimer structures reveals that interactions around R355 and D614 both vary, coupling an important SARS-CoV-2 variant (D614G) to the charge pair examined currently (D398 – R355). Further, and adjacent to the region around D614, interactions surrounding A570 also vary between LA bound and unbound closed S trimer structures (Figure 5). This is of interest since A570D is one of a set of S protein mutations that contribute to the B.1.1.7 lineage of SARS-CoV-2 reported towards the end of 2020 [26]. It has been suggested that other S protein mutations in this variant may alter receptor binding or possibly interaction with antibodies [26]. Thus far A570D has not been included in those discussions. From the current work, since there is a concerted change in structural environments, it is suggested that A570D could alter the balance of closed to open conformation through a similar mechanism to D614G. Although an effect is clearly predicted for A570D, due to the differences in interactions around A570 shown in Figure 5, whether it is as large as that reported for D614G [20], and whether it would tend towards further opening (beyond the D614G level) or revert to more closed (towards the D614 level), is unknown. In the absence of a structure for A570D, this uncertainty is shown schematically in Figure 5B.

In a series of structures of furin cleaved S trimers [25], most of which were released after collation of the trimer dataset, predicted D398 pKas for one extreme (6zgi, all monomers down) are all stabilising, those for all monomers open (7a98, 3 ACE2 bound) are all destabilising. Intermediate combinations of open/closed monomers within a trimer are predicted to have mixed (stabilising, destabilising) pKas. These results, for a set of trimer structures from the same laboratory, that vary from entirely closed to entirely open, are in agreement with calculations for the original trimer dataset, with the consequence being a predicted pH-dependence of the open/closed equilibrium.

Analysis of the effects of mutation to the full set of amino acid alternatives, at RBD locations, recapitulates the divide between RBD subdomains located proximal and distal to the ACE2 binding region [11]. Greater tolerance of mutations is observed, with respect to both ACE2 binding and RBD expression, in the distal subdomain, consistent with the higher B-factors seen in crystal structures. Both R355 and D398, which lie at a junction between these subdomains, are uniquely favoured for expression, and largely favoured for ACE2 binding [11]. This is not surprising if the coupling between R355 and D398 is important for stability. Although D398 is predicted to be destabilised in many of the structures, this is more than compensated by predicted stabilisation of R355, due to the pair interaction. It is interesting that for F464, which occludes both R355 and D398, the two mutations that are tolerated for expression are tyrosine and glycine. Tyrosine is presumably a swap for similar, whereas glycine would increase solvent accessibility around the charge pair and reduce the coupling, as suggested for the RBD of HKU1. In circulating SARS-CoV-2 genomes, mutations of R355, D398, (or F464), are not currently accumulating [27], in line with a functional role. Residue Y423 is positioned to accept or donate a hydrogen bond interaction with D398, so that the buried environment of D398 appears to be compatible with either ionised or protonated sidechain. Tyrosine 423 is also a key residue in the scanning mutation analysis [11].

Cold sensitivity of a spike protein construct is reduced when the protein is engineered to bias towards the closed/down trimer [28]. It is also significantly reduced when the pH is reduced from 7.4 to 6.0, although the molecular origin of this pH effect, and thus whether the D398 – R355 charge pair is involved, is unknown. The scope for a functional role of pH in the viral infection cycle is apparent at multiple points, demonstrated for example in a study of interferon stimulated genes in viral replication [29] that includes endosomal (pH 6.3 to 5.5), and secretory pathway (pH 7 to 5) candidates. It has been reported that evolution of the furin cleavage site at the S1/S2 junction and the D614G mutation act to balance infectivity and stability of SARS-CoV-2 [30]. The current work suggests that the D398 – R355 pair could be another factor in this balance, including a pH-dependent element.

## Conclusions

The expanding dataset of SARS-CoV-2 S protein trimers and fragments has been leveraged to yield pKa predictions that focus on a small number of amino acids, when interrogated for those that show electrostatic frustration. Prominent amongst these is the partially buried salt bridge D398 - R355 in the RBD, which is predicted to contribute a stability difference between open and closed forms, due to opposing burial (destabilising) and charge pair interaction (stabilising) terms. Variation with pH is also predicted, of potential relevance to virus cell entry and exit. The importance of charge coupling between R355 and D398 could be examined with double mutation, simultaneously replacing both ionisable groups. Recent solution of S protein trimer structure with linoleic acid bound shows that monomer – monomer packing and interactions adjacent to the D398 – R355 pair are structurally coupled to those around D614, as well as the environment of A570 (relevant to the B.1.1.7 variant of SARS-CoV-2 that carries the A570D mutation). It is therefore suggested that all of these factors may be linked to the open/closed trimer equilibrium, and therefore potentially to virus infectivity.

## Materials and methods

### Structural datasets

A set of 24 S trimers for SARS-CoV-2 was created from the August 2020 RCSB/PDB, with the S proteins extracted from complexes where necessary: 6vsb, 6vxx, 6vyb, 6xkl, 6z43, 6xm5, 6×6p, 6×29, 6×2c, 6×2a, 7byr, 6zge, 6zgg, 6xcm, 6xs6, 6zp1, 6zp0, 6zoy, 6xr8, 6zox, 6xlu, 6xm4, 6xm3, 6xm0. These are all cryo-EM structures, with resolutions mostly in the range 3 – 4 Å. A recent S trimer with linoleic acid bound was also studied (6zb4).

The RCSB was searched for X-ray crystallographic structures of coronavirus RBD and NTD, with a specific subset collated for SARS-CoV-2 data, in September 2020. These X-ray structures are generally higher resolution than the S trimer cryo-EM structures. Retrieved data for SARS-CoV-2 RBD (PDB and chain) were 6lzgB, 6mojE, 6w41C, 6ylaA, 6yz5E, 6z2mA, 6zczA, 7bwjE, 7c8vB, and chain E of the chimeric 6vw1 structure. For structures of RBDs obtained from human coronaviruses other than SARS-CoV-2, single representatives were taken: 2ghvE (SARS-CoV-1), 4kqzA (MERS), 5kwbA (HKU1), and 6nzkA (HuOC43), where 6nzkA is a cryo-EM structure in the closed form and the others are X-ray crystal structures. A dataset of NTD representative structures was also collected for coronaviruses that infect humans, 6×29A (SARS-CoV-2), 5×4sA (SARS-CoV-1), 5×4rA (MERS), 6nzkA (HuOC43), 5i08C (HKU1). Of these, 6×29, 6nzk, and 5i08 are cryo-EM structures.

### Sequence datasets

In order to visualise amino acid sequence conservation alongside electrostatics calculations, a set of 10 S protein sequences was collected, covering the coronavirus family, with UniProt [31] identifiers and coronavirus sub-family: P0DTC2 (SARS-CoV-2, beta), P59594 (SARS-CoV-1, beta), K9N5Q8 (MERS, beta), A3EXG6 (Bat-HKU9, beta), P11224 (MHV-A59, beta), P11223 (Avian-IBV, gamma), Q91AV1 (Porcine-EDV, alpha), P15423 (Human-229E, alpha), P10033 (Feline-IPV, alpha), and B6VDW0 (Bulbul-HKU11, delta). Sequences were aligned with Clustal Omega [32] at the European Bioinformatics Institute [33], and visualised in ESPript [34]. A conservation score was calculated from the number of occurrences of each SARS-CoV-2 amino acid in the alignment of 10 sequences, colour-coded for display (Figure 1).

### Electrostatics calculations

A model for combined Finite Difference Poisson Boltzmann (FDPB) and Debye-Hückel (DH) continuum electrostatics calculations, termed FD/DH, has been introduced and benchmarked [16]. A feature of this model is the ability to discriminate between largely buried ionisable groups, that may individually contribute substantially to folded state stabilisation or destabilisation (the latter termed here electrostatic frustration), and those groups that are solvent accessible and typically contribute less. The latter includes most salt-bridge interactions, which are folded-state stabilising and are normally plentiful across a protein surface. Further down the scale of interaction strengths are ionisable groups present in a protein but not ordered in a cryo-EM or X-ray structure. In the absence of available coordinates such groups are omitted from calculations, with the assumption that when disordered they are largely solvent accessible with relatively low pKa deviations from normal. Recently a web tool has been reported (www.protein-sol.manchester.ac.uk/pka) [17], that allows users to visualise predictions of ionisable group pKas around specified centres, with the FD/DH method. In the current work, an offline version of the method is used to make pKa predictions for (residue/sidechain group intrinsic pKa) Asp/4.0, Glu/4.4, Lys/10.4, Arg/12.0, and His/6.3 groups, for submitted structures [16], without the requirement for centre and subset specification of the rapid turnaround web tool. Calculations are made for 24 SARS-CoV-2 S trimers, with results visualised as a heat map of predicted pKa deviations, colour-coded as for the web tool, and presented in columns of aligned groups. Where the S protein numbering differs from that of the base structure (6vsb), a register correction is made to maintain the 6vsb numbering scheme. Average predicted ΔpKa is calculated for each ionisable group and placed into a structure file for comparable visualisation (using PyMOL) to that of the web tool (red destabilising, blue stabilising). Swiss PDB Viewer [35] was also used for molecular visualisation, and additionally for alignment of structures.

## Key Points

- Application of pKa calculations across the rapidly increasing dataset of SARS-CoV-2 spike protein trimers and fragments yields a set of consensus predictions that focus on a small number of amino acids.
- A partially buried salt bridge (D398 – R355) in the receptor binding domain is predicted to contribute a stability difference between open and closed forms, which varies in the mild acidic pH region, of relevance to cell entry of the virus and its secretion.
- The salt-bridge lies at a nexus of protein conformational variation, is present in SARS-CoV-2, SARS-CoV-1 and MERS, but either absent or predicted to be less influential in other human coronaviruses.
- A strategy of simultaneous mutation for D398 and R355, and characterisation, would probe the charge pair role in SARS-CoV-2 infectivity.

## Acknowledgements

Staff of the University of Manchester Computational Shared facility are thanked for technical support.

## Data availability statement

The data underlying this article will be shared on reasonable request to the corresponding author.

